# Molecular basis of UV lesion binding and repair inhibition by ETS-family transcription factors

**DOI:** 10.64898/2026.01.13.699325

**Authors:** Smitha Sivapragasam, J. Ross Terrell, Markus W. Germann, Marian F. Laughery, Michaela E. Everly, Shiva P. Adhikari, Patrick J. Hrdlicka, John J. Wyrick, Gregory M. K. Poon

## Abstract

Mutation hotspots in melanoma frequently occur at DNA binding sites of ETS-family transcription factors, as ETS factors stimulate the formation of UV-induced cyclobutane pyrimidine dimers (CPDs) while suppressing repair at ETS-bound DNA sites. To elucidate the molecular mechanism by which ETS factors bind to damaged DNA sites and inhibit repair, we investigated the binding of members from the three major classes of the ETS superfamily (Ets1, ELF1 and PU.1) to cognate DNA containing a *cis-syn* TpT CPD. These site-specific CPDs modulated ETS recognition and repair by a model repair enzyme in a position-dependent manner, with a deaminated CPD located in a damage hotspot in the ETS binding motif consistently stimulating binding and repair inhibition by all three paralogs. Co-crystal structures of PU.1 reveal that CPDs and mismatches are recognized within the framework of canonical ETS/DNA complexes, but with highly differentiated thermodynamic and dynamic properties. Specifically, the CPD-bound complex exhibits unique dynamics that reveal a novel DNA-binding mode as well as inform how ETS domains navigate the DNA conformational landscape to predispose CPD induction. The results offer a molecular rationale for how ETS factors induce mutation hotspots in skin cancers and other UV-exposed tissues by promoting CPD induction and inhibiting repair.

**KEY POINTS:**

1. ETS-family proteins from all major classes bind CPD-containing DNA sites.
2. Engagement of CPD by ETS proteins inhibits DNA repair in a position-dependent manner.
3. Structure and thermodynamics of ETS binding to damaged DNA differ from undamaged substrates.

## INTRODUCTION

Ultraviolet (UV) light is the primary etiological agent for skin cancers such as melanoma by inducing mutagenic damage to DNA (1,2). The most common type of UV photodamage is the cyclobutane pyrimidine dimer (CPD), which is formed by a [2+2] cycloaddition reaction between the C5-C6 double bonds of neighboring pyrimidine bases (3). CPDs are repaired by the nucleotide excision repair (NER) pathway (4), which plays a critical role in preventing carcinogenic UV-induced mutations. Xeroderma pigmentosum patients with inherited defects in NER have >1000-fold increased risk of skin cancer (5). Previous genome-wide surveys of NER activity in human cells using the excision repair-sequencing (XR-seq) method have indicated that the rate of CPD repair is remarkably variable across the genome, with slow repair associated with locations where access to DNA damage is restricted by chromatin architecture or DNA-bound proteins (6,7). Since the efficiency of repair significantly impacts UV-induced mutation rates, it is important to elucidate the molecular mechanisms that regulate lesion accessibility and repair activity.

Somatic mutations in melanoma are highly enriched at DNA sites bound by transcription factor (TF) proteins (8–10). Some of these mutation hotspots have been identified as non-coding driver mutations, including, for example, recurrent mutations that create binding sites for E26 transformation-specific (ETS) family TFs in the promoter of the telomerase reverse transcriptase (*TERT*) gene (11,12). However, there are recurrent somatic mutations at many other TF binding sites in melanomas that are not linked to carcinogenesis (13). Instead, analysis of genome-wide repair of CPD lesions in yeast and human cells indicates that DNA binding by certain classes of TFs may promote UV mutagenesis by inhibiting NER (8,14,15). For example, the CCCTC-binding factor (CTCF) inhibits the repair of CPDs at CTCF binding sites across the genome, which correlate with elevated somatic mutation rates in melanoma at these binding sites (8,16,17). In parallel, we have shown that purified CTCF protein binds with only slightly lower affinity to a CPD-containing DNA binding site *in vitro*, and inhibits CPD cleavage by T4 endonuclease V, a model CPD repair enzyme (18). Other transcription factors, however, vary in their ability to tolerate CPD lesions in their cognate binding sites. Early studies identified several prominent examples (p53, AP-1, NF-Y, E2F4 and NFκB) with significantly decreased binding affinities to site-specific CPDs (19). Recent high-throughput characterization has indicated that impaired TF binding to damaged binding sites is widespread (20), and that many TFs, including the ETS-family members ETS1 and ELK1, show altered DNA binding specificity for UV damaged DNA. However, the molecular mechanisms that regulate TF binding to damaged DNA sites remain unclear.

DNA binding sites of ETS family TFs are among the most highly mutated non-coding elements in melanoma (9,21–27). These mutation hotspots occur at least in part due to induction of CPD formation at specific locations in ETS-bound DNA sites (21,22,26–28), as ETS TF binding alters the DNA structure so that C5-C6 double bonds of neighboring pyrimidine bases have distances and torsion angles favorable to CPD formation (21). One of these damage and mutation hotspots occurs at the central TpC dipyrimidine step in the central 5’-(A/T)<u>TC</u>C-3’ ETS binding consensus. Notably, a recent genome-wide analysis indicates the TpC CPDs at this location in the ETS motif rapidly deaminate to uracil (i.e., to a deaminated TpU CPD) and are repaired slowly by the NER pathway *in vivo* (14). A previous study indicates that the Class I ETS TF (i.e., ETS1) may inhibit repair by blocking accessibility of CPD lesions to the NER damage sensor UV-DDB, at least *in vitro* (20). However, the molecular mechanism by which ETS factors accommodate a CPD lesion in their DNA binding site is unclear. This is particularly true for Class II and III ETS family members, which comprise more than half of the 28 human ETS paralogs (29) and are also normally (30) as well as pathologically (31) expressed in epidermal and dermal tissues with exposure to incident UV radiation. While the structures of CPD-containing DNA alone (32) or in complex with DNA repair factors such as photolyases (33,34), UV-DDB (35), or xeroderma pigmentosum C (36) has been previously characterized, how ETS proteins or other TFs accommodate a CPD lesion in their DNA binding site is not known.

To this end, our objectives in this work were three-fold. First, we determined the DNA-binding properties of archetypal members of three ETS family classes (I-III), in order to test whether binding and repair inhibition at CPD-bearing DNA sites represents a class property of the ETS family. Second, we solved high-resolution co-crystal structures to gain insight into the impact of CPD and downstream Watson-Crick mismatches (due to CPD deamination) on ETS/DNA complexes. Third, we dissected the structural thermodynamics of ETS binding to these DNA substrates to probe the energetic and dynamic basis of CPD formation and recognition by ETS proteins. The results provide a deep characterization of how a bulky helix-distorting lesion is accommodated within ETS TF binding sites and interferes with subsequent repair, thereby promoting mutation hotspots at ETS binding sites in skin cancers.

## MATERIALS AND METHODS

### Nucleic acids

Synthesis of the CPD oligo is detailed in *Supplemental Methods*. Non-CPD oligos were purchased from Integrated DNA Technologies (Midland, IA). Dried oligos were dissolved in aqueous 1.0 M NaCl and desalted on G-25 resin (HiTrap Desalting columns; Cytiva) in water under the control of an ÄKTA start (Cytiva) system. The salt-free high-MW fraction was lyophilized and redissolved in 30 mM HEPES, pH 7.5 containing 100 mM potassium acetate. Duplexes were annealed from equimolar amounts of the desired complementary sequences (in ∼200 µL volume) by heating to 90°C followed by passive cooling to room temperature. Duplex concentrations were determined by UV absorption at 260 nm using the nearest-neighbor extinction coefficients for the WC sequence (37).

### Protein purification

pET28b-based (Novagen) constructs encompassing the C-terminal ETS domains of murine Ets-1 (residues 280 to 440, a kind gift from Dr. Lawrence McIntosh) or human PU.1 (residues 165 to 270) were transformed into BL21(DE3)pLysS *Escherichia coli* (Invitrogen). Residues 161 to 340 of human ELF1 encompassing its internal ETS domain were cloned into pCDF-1b (Novagen) and similarly transformed. Cultures in LB medium were induced with 0.5 mM isopropyl β-D-1-thiogalactopyranoside at OD_600_ of 0.6 for 4 hours at 30°C. Harvested cells were re-suspended in Buffer H (10 mM HEPES, pH 7.4, with 0.5 M NaCl and 0.5 mM Tris(2-carboxyethyl)phosphine) containing 1 mM phenylmethylsulfonyl fluoride) and lysed by sonication. The lysate was cleared by centrifugation and loaded onto a HiTrap SP HP column (Cytiva) equilibrated with Buffer H. After washing, the protein was eluted along a linear NaCl gradient under the control of a Bio-Rad NGC instrument. Samples for co-crystallization were concentrated in Amicon Ultra-15 centrifugal filters (10k MWCO) and then polished and exchanged into Buffer H on a HiLoad 16/600 Superdex 75 column (Cytiva). Protein concentration was determined by UV absorption at 280 nm based on an extinction coefficient (in M^-1^ cm^-1^) of 39,880 for Ets1, 28,420 for ELF1 and 22,460 for PU.1.

### Electrophoretic mobility shift assays

For DNA-protein binding assays, 6-FAM-labeled duplex oligonucleotides (200 nM) harboring native and variously modified ETS-binding sequences of the *DPH3* promoter were titrated with ETS domain in 25 mM Tris HCl (pH 7.9) containing 0.5 mM EDTA, 6 mM MgCl_2_, 60 mM KCl, 10% glycerol, 10 mM DTT, and 0.12% w/v of bovine serum albumin (BSA) in a total volume of 50 µL. The DNA-protein mixture was incubated on ice for 45 minutes and 5 µL of the mix was loaded onto an 8% native polyacrylamide gel. Electrophoresis was performed at 30 V/cm for 30 min. Gels were imaged using a Typhoon FLA700 scanner (Cytiva). Bands were quantified using ImageQuant software (Cytiva).

### T4 pyrimidine DNA glycosylase (PDG) inhibition assays

From the gel mobility shift samples, 10 µL of DNA (200 nM) and ETS domains (at concentrations corresponding to ∼95-100% saturation) were treated with 10 U of T4 PDG (New England Biolab) at 37°C. Samples were collected after 15 and 30 min and quenched with 2 µL of 1 M NaOH and heated at 95°C for 15 min. Formamide was added to 70% v/v and the samples were further heated at 95°C for 15 minutes. From each sample, 10 µL was loaded onto a 12% polyacrylamide gel containing 8 M urea in 0.5 × TBE buffer. Electrophoresis was carried out at 30 V/cm at 60°C for 20-25 minutes. Gels were imaged using a Typhoon FLA7000 imager (Cytiva) quantified using ImageQuant software (Cytiva).

### X-ray crystallography

Purified protein was mixed with target DNA at 1:1 molar ratio in Buffer H to yield a complex concentration of 200 µM. Co-crystals were grown for 5 days by vapor diffusion at 293 K in a 2 µL hanging drop comprised of a 1:1 mixture of protein:DNA complex with mother liquor containing 100 mM sodium acetate, pH 4.6, and 2% PEG 3350. Prior to freezing, 2 µL of a cryoprotectant solution containing 100 mM sodium acetate, 2% PEG 3350, and 20% glycerol was laid on top of the hanging drop and the well closed for 1 h of incubation (4 µL total volume, 10% glycerol concentration). After 1 h, crystals were transferred to the above 20% glycerol solution prior to freezing. X-ray diffraction data sets were collected at the National Synchrotron Light Source II at Brookhaven National Laboratory (Upton, NY).

The diffraction data was processed using the XDS package (38) and scaled using Aimless in the CCP4 package (39). Molecular replacement was performed using a PU.1 co-crystal complex (PDB: 8E3K) as the search coordinates in the PHENIX suite (40) via the maximum-likelihood procedures in PHASER. Rounds of refinement were then carried out using phenix.refine (40) followed by model building in Coot (41). Refinement statistics and information regarding specific beamlines, detectors, collection wavelengths, and oscillation angles are given in **Table S1** (Supporting Information).

### Fluorescence polarization titrations

A Cy3-labeled DNA probe (42) encoding a high-affinity sequence for PU.1 (5’-AGCGGAAGTG-3’; 0.5 nM) was incubated with graded concentrations of protein and unlabeled DNA to equilibrium (overnight at 25°C) in 10 mM TrisHCl pH 7.5 containing 0.01% BSA and 0.15 M NaCl. Steady-state fluorescence anisotropies were measured at 595/30 nm in a Molecular Dynamics Paradigm micro-plate reader with 530/30 nm excitation.

Observed anisotropies, computed from buffer-subtracted plane-polarized fluorescence, were reported as means ± S.D. of three or more experiments:

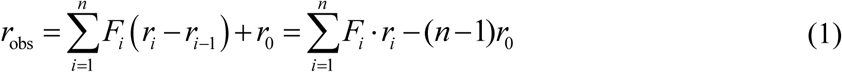

where *r*_0_ represents the anisotropy of the free DNA probe and *r_i_* refers to the *i*-th bound species.

*F* is the fractional bound probe and was analyzed as a function of total protein concentration as follows.

### Model-dependent analysis of fluorescence polarization titrations

As previously established by calorimetric (43) and diffusion measurements (44) in solution, PU.1 (P) binding to a single cognate DNA site (D of species *j*) sequentially with negative cooperativity:

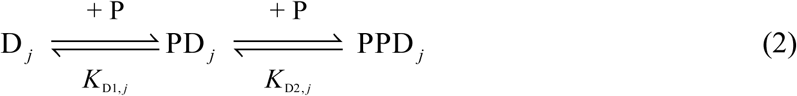

where *^K^*_D1_ and *^K^*_D2_ are the equilibrium dissociation constants describing the 1:1 and 2:1 PU.1/DNA complexes ( PD *_j_* and PPD *_j_* ):

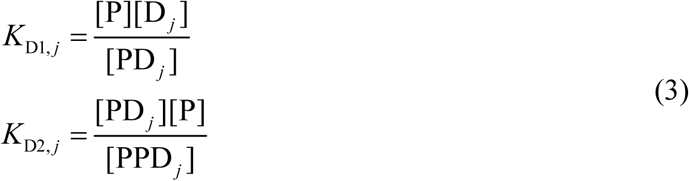

The concentrations of the various species were solved by numerical root finding, as detailed in (42), to compute the fractional bound probe (taken as species D_1_ ), *F*_1_ :

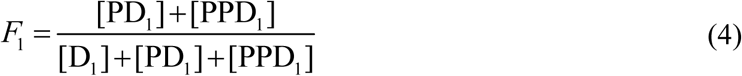

This approach accounts for mutual depletion among all species, a detailed exposition of which is found in (45), and yields absolute binding affinities (without reliance on IC_50_ values). To reduce the degrees of freedom in competition (*j* = 2) experiments, protein was deployed at sub-saturating concentrations with respect to the probe. Nonlinear least square regression of the system of equations consisting of Eqs. (1) and (3) was performed using Origin software (Northampton, MA).

### Isothermal titration calorimetry

Purified DNA and ETS domain were extensively co-dialyzed in 10 mM NaH_2_PO_4_/Na_2_HPO_4_ at pH 7.4 with 0.15 M NaCl. Dialysate was used in all dilution and rinsing procedures. DNA was loaded into the cell (*V* = 1.4 mL) at a nominal concentration of 10 µM and protein at 200 µM in the syringe (300 µL) of a VP-ITC instrument (Malvern). DNA (or buffer) was titrated, with constant stirring, with an initial 2.0-µL injection, followed by thirty 9.0-µL injections of protein spaced at 300-s intervals at 25°C. After baseline subtraction, heat peaks were integrated and expressed as 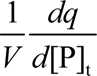 *i.e.*, heat per mol of injected titrant as a function of the titrant:titrate molar ratio *X* (with volume corrections made by the acquisition software).

Experimental per-injection heats were modeled according to an initial value problem as described previously (46):

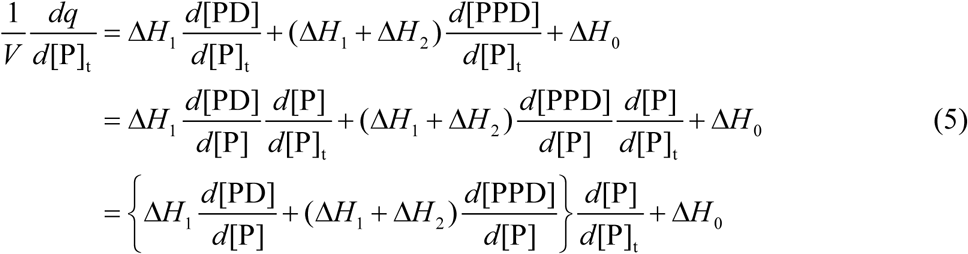

where the indices 1 and 2 on the enthalpy changes refer to the sequential species PD and PPD, and 0 corresponds to the baseline corresponding to the residual heat of protein dilution. The three derivatives that satisfy the sequential binding model as embodied by Eq. (2) are:

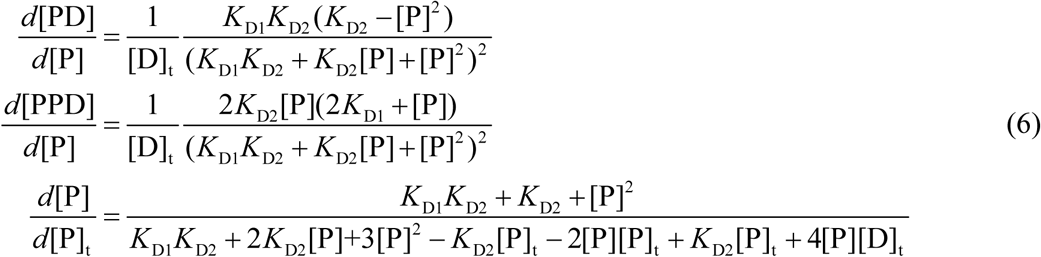

The macroscopic equilibrium constants *K*_D1_ and *K*_D2_ are as defined in Eq. (3). [P]_t_ is transformed from experimental independent variable *X* as follows:

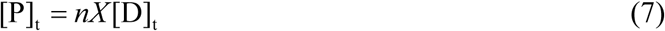

where the subscript “t” refers to the total concentrations and *n* is a correction factor for capturing concentration deviations from nominal values. Typically, 0.9 < *n* < 1.1. Numerical solution of the initial value problem by Runge-Kutta methods was implemented using the d02pvc/d02pdc functions from the NAG C (Numerical Algorithms Group, Oxford, UK) library, as previously described (46), and nonlinear regression was performed using Origin. The reported enthalpy change for the 1:1 PU.1/DNA complex Δ*H*_1_ (corresponding to the co-crystal structures) was taken as the fitted difference from the first transition to the final baseline at high-excess of titrant, adjusted to the experimental heats from protein-into-buffer titrations.

### Molecular dynamics simulations

Explicit-solvent simulations were performed with the Amber14SB force field with the parmbsc1 modifications for DNA (47) in the GROMACS 2025.*x* environment. DNA duplexes were constructed in B-form geometry using 3DNA. The co-crystal structure (8E3K) was used as initial coordinates of the wildtype PU.1/DNA complex. Each system was set up in dodecahedral boxes at least 1.0 nm wider than the longest dimension of the solute, solvated with TIP3P water, and neutralized with Na^+^ and Cl^-^ to 0.15 M. Electrostatic interactions were treated by particle-mesh Ewald summation with a 1 nm distance cutoff. All simulations were carried out at 298 K with velocity rescaling (48). A timestep of 2 fs was used and H-bonds were constrained using LINCS. After the structures were energy-minimized by steepest descent, the *NVT* ensemble was equilibrated for 1 ns to thermalize the system, followed by another 1 ns of equilibration of the *NPT* ensemble at 1 bar (C-rescale barostat). The final *NPT* ensemble was simulated without harmonic restraints (specified as follows), recording coordinates every 1 ps. In all simulations, distance restraints were placed in the terminal DNA base pair to maintain the center-of-mass (COM) of the H-bonding heavy atoms at the energy-minimized distances (force constant *k*_B_ = 10^6^ kJ mol^-1^ nm^-2^). Restraints (when used) in CY base steps consisted of a distance restraint between the corresponding C5 and C6 atoms (target value 1.5 Å and *k*_B_ = 6 × 10^4^ kJ mol^-1^ nm^-2^), and/or a dihedral restraint defined by the C5-C6 vectors (C5-C6-C6*-C5*; target value 16° with *k*_B_ = 2 × 10^3^ kJ mol^-1^ rad^-2^, equivalencing 1 unit distance with 2π radians). Convergence of the trajectories was checked by RMSD of the energy-minimized structures, after corrections for periodic boundary conditions (PBC). Triplicate production runs were carried out using different random seeds in the velocity distribution. Analysis of trajectories was performed using tools provided by the GROMACS suite, with entropy estimation further elaborated as follows.

Absolute conformational entropies were estimated according to Schlitter (49). PBC-corrected trajectories were further fitted to remove rigid-body rotational and translational degrees of freedom. The final 5 ns of each trajectory was used as input to generate a covariance matrix of atomic fluctuations. For DNA, the trajectory was fitted to the internal 10 base pairs (to minimize artefacts from terminal fraying); for proteins, the trajectory was fitted to the backbone heavy atoms. The resulting eigenvalues were then used to compute the Schlitter entropy at 298 K.

## RESULTS

### Paralogs from all three major ETS classes can bind a CPD-containing ETS-binding motif

To characterize how CPD lesions impact binding of ETS family TFs, we synthesized double-stranded oligonucleotides containing site-specific CPD lesions in a model ETS binding site. This binding site was derived from the *DPH3*/*OXNAD1* bidirectional promoter (henceforth *DPH3* promoter), which regulates the *diphthamide biosynthesis 3* (*DPH3*) and *oxidoreductase NAD-binding domain containing 1* (*OXNAD1*) genes. This site was chosen because it is highly mutated in sequenced melanoma tumors due to ETS-induced CPD formation hotspots (9,21,26,28,50). We synthesized oligonucleotides with a *cis-syn* TpT CPD starting at positions -2 (**CPD_-2_**) and -1 (**CPD_-1_**) relative to the ETS motif midpoint (Fig. 1A). Position -2 corresponds to a CPD forming between two conserved thymine residues in the ETS binding consensus, while position -1 models a deaminated CPD (*i.e.,* TpU) initially forming at a TpC base step (Fig. 1A). Previous studies indicate that ETS TF binding does not induce CPD formation at position -2, but induces a CPD hotspot at position - 1 (21,22,27). Importantly, TpC CPDs at position -1 have been shown to rapidly deaminate (i.e., to TpU) in cells relative to other positions in the ETS binding motif (14), which may play a role in mutagenesis.

**Figure 1.**
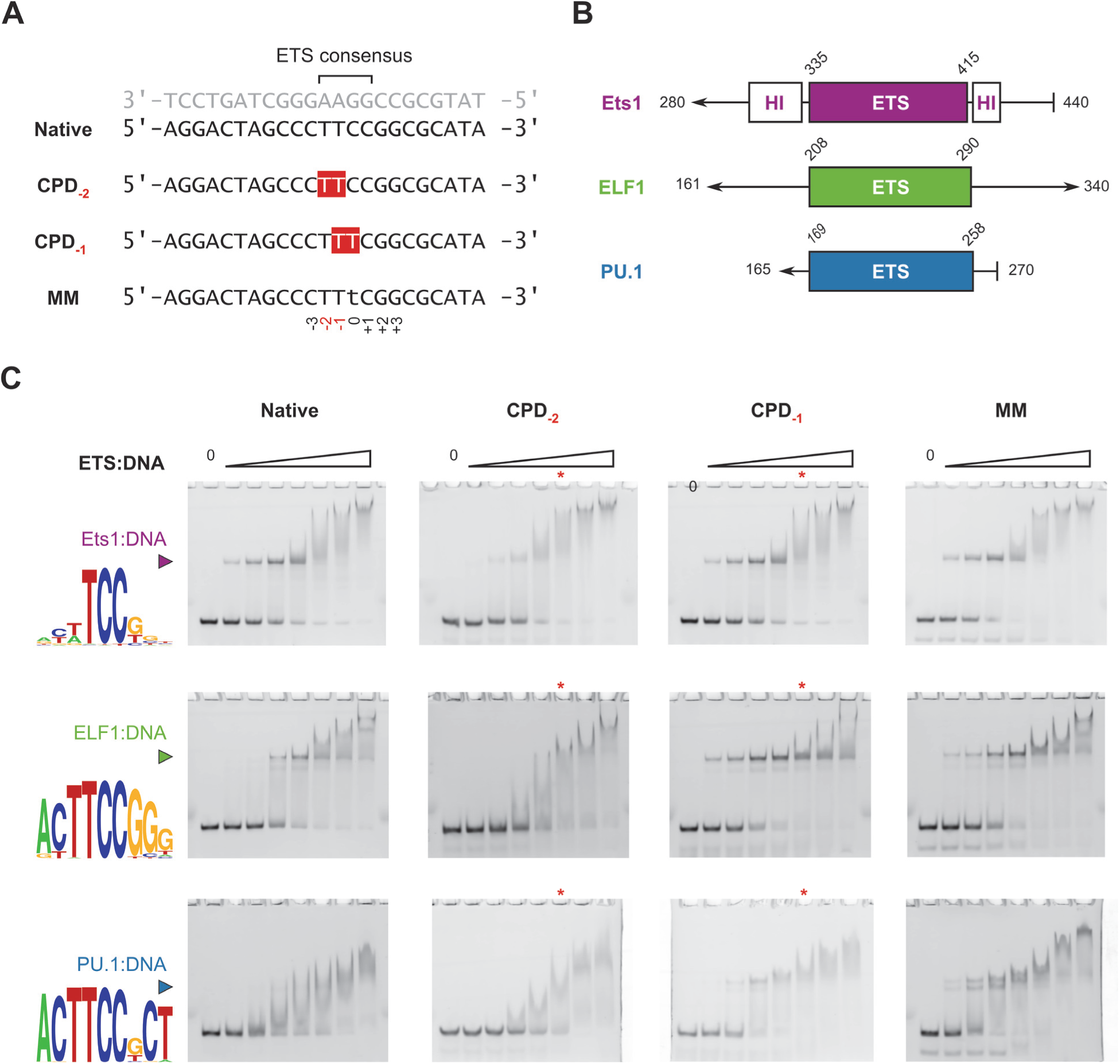
Variation in binding affinity of ETS factors to CPD-bearing and mismatched DNA within the ETS motif in the *DPH3* promoter. **A**, Sequences of the DNA targets studied. For concision, only the pyrimidine-rich sequences of the target duplexes are listed explicitly. **B**, Schematics of the domain structure of murine ETS1 (Class I) and human ELF1 (Class II) and PU.1 (Class III). **C.** Representative gel electrophoretic mobility titrations of the DNA targets by the three ETS representatives. Colored *triangles* denote the mobility of the putative 1:1 ETS/DNA complex. Gel images are rendered as digitized without adjustments. DNA logos for each protein were taken from the CIS-BP database (52). The numerical ETS:DNA ratios are as follows: Ets1: {0.5, 1, 2, 6, 11, 23, 46}; ELF1: {1, 2.5, 5, 9, 17, 26, 52}; PU.1: {0.5, 1.5, 2.5, 5, 13, 26, 39}. The red asterisks refer to the ratios used in the T4 PDG cleavage assay (see Figure 2).

To evaluate the potential for ETS paralogs from all three classes to engage these CPD-containing DNA sites and inhibit their repair, we compared the binding of the purified ETS domains of ETS1 (Class I), ELF1 (Class II), and PU.1 (Class III) to duplex oligonucleotides encoding the native and CPD-modified or Watson-Crick (WC) mismatched *DPH3* sequences [**Figure 1A and B**]. These paralogs were selected based on their representativeness of their respective class and (to maximize the robustness of the findings) the diversity of their domain architecture (29). Since the TpT CPD at position -1 results in a T:G mismatch, we also analyzed ETS paralog binding to an oligonucleotide with a Watson-Crick T:G mismatch at the same location (**MM** control; *c.f.*, Figure 1A).

Titration of the native *DPH3* promoter with the ETS domains of the three paralogs gave rise to a single discrete bound species by electrophoretic mobility shift around the fixed DNA concentration (0.2 µM), corresponding to a 1:1 ETS/*DPH3* complex [**Figure 1C**]. The relative mobility of this species reflected the size of the recombinant constructs (Figure 1A). Addition of excess ETS protein led to secondary high-stoichiometric or nonspecific complexes with greatly reduced electrophoretic mobility. On this basis, the presence of a *cis*-*syn* dithymidyl (TpT) CPD dimer perturbed the apparent affinity for the altered motif in a position-dependent manner. More precisely, replacement of the TT dinucleotide in the 5’-<u>TT</u>CC-3’ consensus by a CPD (**CPD_-2_**) reduced the affinity of the 1:1 complex. However, a centered CPD in the consensus (**CPD_-1_**, 5’-T<u>TT</u>C-3’), corresponding to the most frequently damaged position in the ETS-bound motif in UV-irradiated fibroblasts (21), did not decrease the affinity of the 1:1 complex, nor did the isomeric TT WC-mismatch (**MM**).

Under the identical conditions of the experiment, ELF1 and PU.1 bound the native DPH3 promoter significantly more weakly than Ets1. This observation could be expected based on the differences in sequence selectivity, specifically the positions flanking the 5’-TTCC-3’ consensus (51). Nevertheless, the presence of CPD altered the apparent affinities of the two paralogs in a similarly position-dependent fashion as Ets1. While **CPD_-2_** further impaired binding by ELF1 and PU.1, the proteins bound **CPD_-1_** markedly better and to a similar extent as the non-CPD-containing mismatch **MM**. In particular, the Class III representative PU.1, which strongly prefers pyrimidines at the +2 and +3 positions (Figure 1B), was strongly disfavored by the native *DPH* promoter, but bound **CPD_-2_**and **MM** with apparently similar affinity as ETS1 and ELF1.

In summary, the presence of CPD significantly modifies the apparent affinity of ETS proteins from all three classes. Importantly, the most frequent ETS-associated CPD, as embodied by **CPD_-1_**, enhances site-specific binding by ETS factors generally, including those (such as PU.1) which binds the native sequence with much lower affinity. The similar binding profiles for **CPD_-1_**and **MM** further show that the enhanced affinity towards ETS factors is maintained following rapid deamination of the TpC CPD formed in the 5’-T<u>TC</u>C-3’ consensus to TpU/TpT.

### ETS TF binding inhibits cleavage of a CPD lesion by a model repair enzyme

The position-dependent effects of CPD binding by ETS-family relatives prompted us to ask whether these perturbations could impact the accessibility of the lesion to repair. To test this possibility, we used the model repair enzyme T4 pyrimidine dimer glycosylase (T4 PDG, also known as T4 endonuclease V), which specifically cleaves CPDs, as a biochemical model to probe the competitive inhibition of DNA repair by ETS proteins. The CPD substrates, alone or in the presence of excess ETS proteins sufficient to achieve ∼95-100% saturation, were treated at 37°C with 5 U T4 PDG, sufficient to reach equilibrium CPD cleavage by 15 min. The timescale of the assay was >40-fold the residence time required for cleavage based on reported values (*k*_cat_ > 0.05 s^-1^) of T4 PDG catalytic turnover (53).

From the gel electrophoretic shift data, most of the substrate was bound or supershifted by ETS protein at the concentrations used in the T4 PDG assay (lanes marked with asterisks in Figure 1C). However, assays of T4 PDG activity showed that cleavage of **CPD_-2_** alone was essentially complete by 15 min and was quantitatively unaffected by any of the ETS proteins [**Figure 2**]. In contrast, for **CPD_-1_**, cleavage of DNA alone reached ∼80% completion but, more importantly, was significantly inhibited by each of the ETS paralogs (*p* < 0.001, Figure 2). Notably, the same concentration of ETS proteins that inhibited repair of **CPD_-1_**had no effect on repair of **CPD_-2_**. Thus, biophysical differences in CPD engagement by ETS proteins are reflected in biochemical inhibition of the model DNA-repair enzyme T4 PDG, consistent with dynamic competition between ETS and T4 PDG for CPD substrates. The data therefore indicate that ETS TF binding to CPD-containing binding sites can inhibit accessibility of the lesion to repair factors.

**Figure 2.**
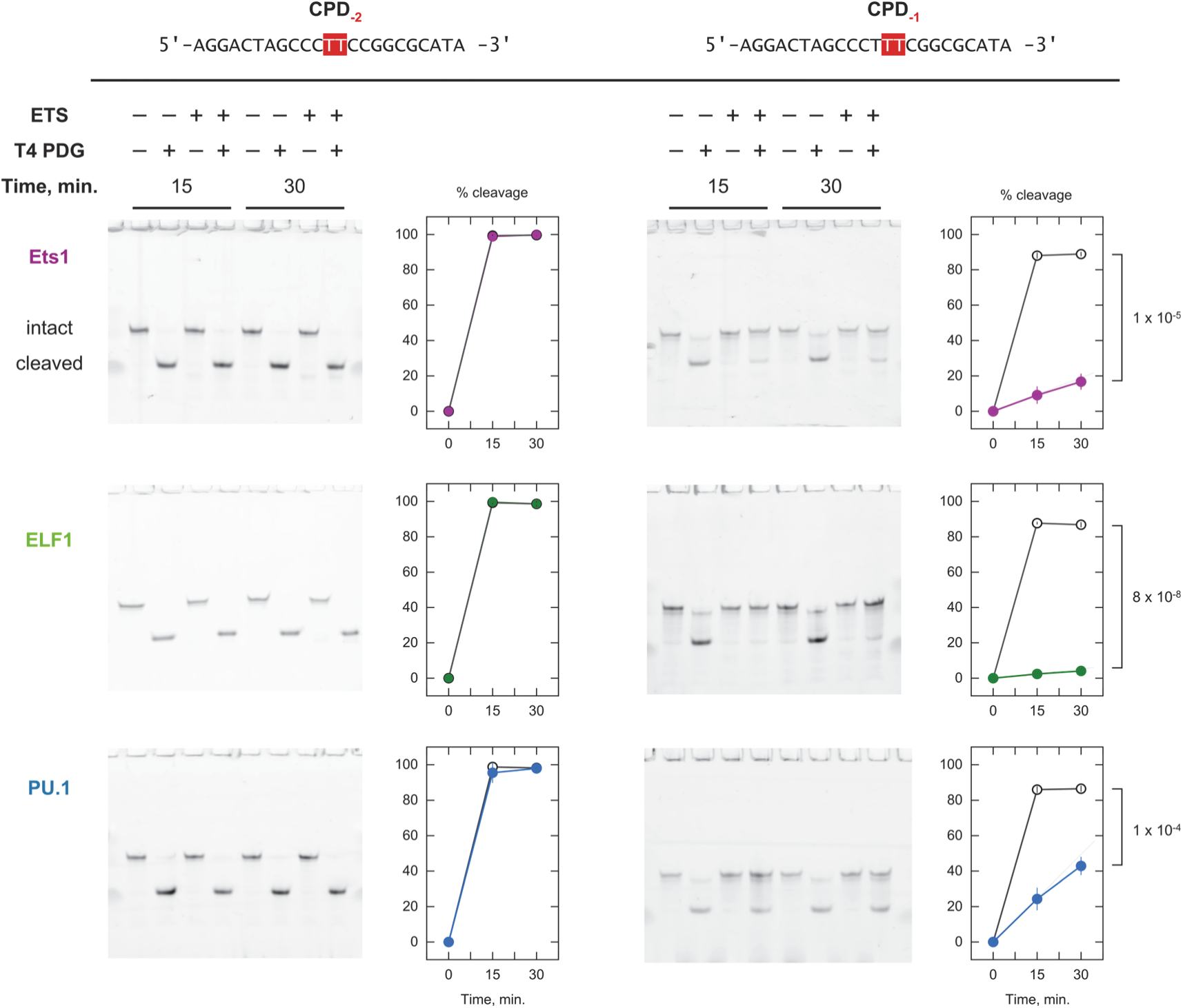
T4 PDG activity at CPD substrates is sensitive to DNA sequence context and ETS binding. Shown are representative denaturing polyacrylamide gels of untreated and T4 PDG-cleaved CPD-bearing substrates sampled at the indicated time points. Fractional cleavage of the substrates under each ETS/DNA combination and condition (filled and open symbols for with and without protein) was analyzed as means ± S.D. from triplicate experiments. Cleavage of CPD_-2_ is quantitatively similar with and without protein at corresponding time points. Gel images are rendered as digitized without adjustments. *p*-values from a two-tailed *t*-test for the difference at 30 min are given for the CPD_-1_ sequence.

### Structural bases of ETS interactions with damaged and mismatched DNA

Co-crystal structures of members across the ETS superfamily have been reported since the PU.1/DNA complex in 1996 (54). These structures demonstrate strong conservation in tertiary structure as well as key DNA contacts by signature residues, principally two arginine sidechains in the recognition helix (H3) that engage the 5’-GGA(A/T)-3’ consensus. Even in complex sequence contexts (55,56) comprising multiple copies of consensus, ETS domains adopt quaternary structures that conserve the domain architecture of each subunit by swapping corresponding secondary elements (57). This structural fidelity is not shared by the DNA-binding domains of other transcription factor families in the helix-turn-helix superfamily, for example the homeodomains and POU family (58). Here, we wished to gain structural and biophysical insight into the recognition of CPD and mismatched consensus DNA by comparing co-crystal structures of DNA bound by ETS factors. Although the structural homology of ETS/DNA complexes at the domain level is a class property of the ETS family, these complexes co-crystallize with diverse degrees of symmetry. The sensitivity of DNA structure *in crystallo* to packing effects imposes a need for ETS/DNA complexes with equivalent crystal packing. To this end, we selected PU.1 which co-crystallizes with a broad range of DNA sequences in an identical space group with the additional benefits of containing a single complex per asymmetric unit and favorable diffraction characteristics (59). The high-affinity PU.1/DNA complex (PDB: 8E3K, herein referred to as **WC**) has among the highest resolutions among deposited ETS co-crystal structures. PU.1 is therefore a useful experimental model for comparative analysis of altered interactions and dynamics induced by structural perturbations such as CPD and WC mismatches.

Co-crystallization of the ETS domain of PU.1 (human residues 165 to 270; ΔN165) with WC variants harboring a (TT-)CPD at the center of the 5’-T<u>TC</u>C-3’ (**CPD**; PDB 9YNZ) or the mismatch (**MM**; PDB 9OA4) 5’-T<u>TT</u>C-3’ yielded complexes [**Figure 3A and B**] packing in the identical space group (P 1 2_1_ 1) as the WC complex. In all the complexes, contacts from the ETS domain were made predominantly in the 5’-GGAA-3’ strand, while positions along the 5’-TTCC-3’ strand corresponding to the CPD and mismatched pyrimidines were not directly contacted by the protein [**Figure 3C**]. The purine strand in **CPD** showed similar contacts with the ETS domain, adjusted for resolution, as **WC**. In the **MM** structure, the wobble-paired guanine (with thymine) is shifted away from the protein, and contact was maintained through an extended sidechain conformation by R230.

**Figure 3.**
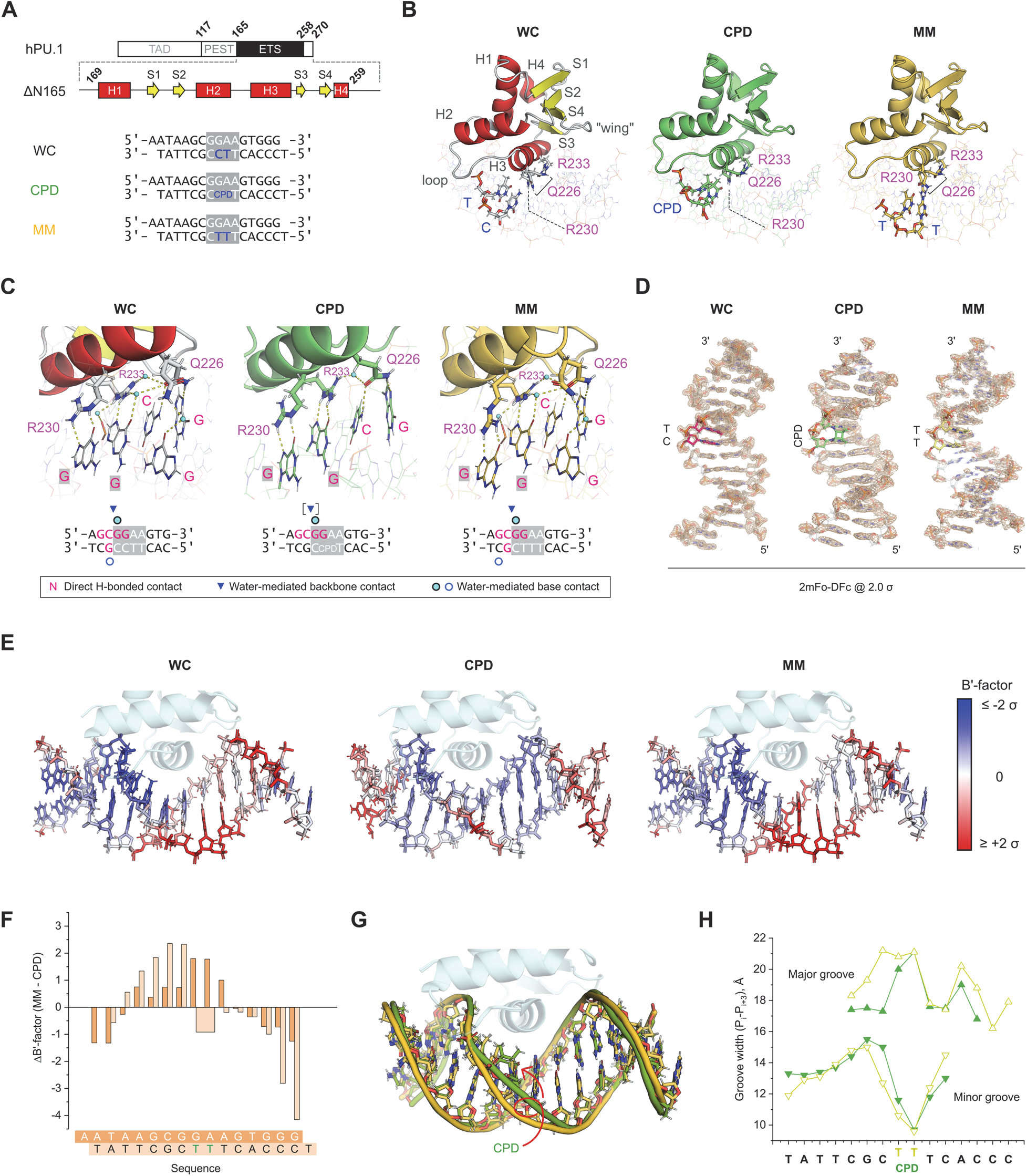
Intermediates of dipyrimidine modifications in complex with the ETS-family transcription factor PU.1. **A**, Schematic of the ETS domain of human PU.1 (residues 165-272; denoted ΔN165) and key DNA oligonucleotides used in the co-crystallization experiments. **B**, Refined co-crystal structures. **C**, Contacts made by the key interfacial residues Q226, R230, and R233. Dashes denote H-bonds within a distance cutoff of 3.5 Å between heteroatoms. The absence of ordered hydration in the CPD-containing structure may be related in part to resolution (1.9 Å), so judgement on water-mediated contacts are reserved (square brackets). **D**, Comparison of 2mFo-DFc maps at a common relative cutoff of 2.0 σ. The variable dinucleotide steps are rendered in color. **E**, B’-factors of each bound DNA mapped to the modeled structures. Two standard deviations in B’-factor (calibrated internally to each structure) are rendered the resolve the implied dynamic distribution among the internal ETS-bound base pairs. **F**, Per-residue ΔB’-factors for heavy atoms between the ETS-bound mismatched and CPD-bearing DNA highlight major differences in implied DNA dynamics between the CPD and MM complexes. **G**, DNA structures of the CPD (green) and MM (gold) complexes, aligned to the bound protein. **H**, Major and minor groove widths, taken as the distance between P atoms of the *i*^th^ and *i*+3^th^ base.

Over the whole protein-bound DNA, the bound CPD showed more distinct differences from the parent WC and mismatched TT sequences [**Figure 3D**]. The CPD lesion exhibited low electron density (2mFo-DFc) while coverage elsewhere was comparable with WC sequence. We considered the possibility of photoreversion by the (TpT) CPD into TT, effectively converting **CPD** to **MM**, during handling. This possibility was discounted by the qualitatively distinct protein/DNA interfaces of **CPD** and **MM** (Figure 3C), as well as the 2mFo-DFc coverage of the MM complex showing reduced density at positions *flanking* the TT base step, in contrast with **CPD** (Figure 3D). To test the robustness of the distinct 2mFo-DFc map in **MM**, we solved a second mismatched structure in which inosine replaced guanine in the G:T base pair and made the same wobble contacts but with a weaker stack. The resultant complex (**I-MM**; PDB 9OB0), which crystallized to an equivalent asymmetric unit [**Figure S1A**], showed a similar 2mFo-DFc map as **MM** [**Figure S1B**].

To directly establish that the CPD sequence was intact, we probed its structure by solution ^1^H NMR, focusing on the imino-^1^H fingerprint region. When annealed with the same complementary strand as the native sequence, the CPD duplex showed the full complement of imino (for G and T bases) chemical shifts. Assigned peaks at and adjacent to the CPD were distinct in comparison with MM and unmodified DNA [**Figure S2B**]. Independently, we probed the CPD enzymatically with T4 PDG and observed quantitative cleavage of a ligated duplex product [**Figure S2C to E**]. The CPD duplex was therefore intact at least up to crystallization.

Based on the evidence, two scenarios present themselves to account for the low bridging density of the bound CPD. The CPD might have been split by synchrotron radiation, in which case the diffraction captured a conformational state prior to relaxation to the MM conformation. The second possibility, not exclusive of the first, is that the 2mFo-DFc maps reflected in part molecular dynamics along the bound DNA. With these considerations, we compared the (*z*-normalized) B’-factors, which quantify isotropic dynamics in the crystal, for the ETS-bound WC, CPD, and MM sequences. Except for a localized hotspot adjacent to the TpT dimer, the DNA dynamics in **CPD** are distributed relatively evenly around their average along the DNA-contacting segment of the DNA duplex [**Figure 3E**]. This contrasts with the biased distribution in **MM** [**Figure 3F**], which is similar to **WC** and other reported non-CPD PU.1/DNA complexes. Structurally, the reduced base-base distance imposed by the TpT dimer disturbs the local helical conformation [**Figure 3G**], such that the major groove is locally narrowed and the minor groove enlarged just upstream of the CPD [**Figure 3H**]. The ETS-bound CPD therefore exhibits distinct structural and dynamic properties as manifest *in crystallo* by way of the distribution of electron density, implied dynamics, and helical conformation.

### ETS interactions engage CPD and mismatched DNA in a thermodynamically distinct binding mode from WC-matched targets

The electrophoretic mobility shift data showed that a TpT CPD centered at the ETS consensus motif (5’-T<u>TT</u>C-3’; CPD-1) was recognized by ETS proteins, including PU.1, on par with the native sequence in the *DPH3* promoter (Figure 1C). To test whether the sequence context of the co-crystal structures reflect the same behavior, we measured the binding affinity of PU.1 for these targets in solution using a fluorescence polarization assay [**Figure 4A**]. Relative to WC, the affinity for CPD was equivalent within experimental uncertainty, while MM was ∼10-fold weaker. We further probed the role of the wobble G:T base pair in the MM using the I-MM sequence. In a Watson-Crick G⫶C context, G>I substitution reduces PU.1-binding affinity due to an uncompensated increase in the local conformational entropy of the wobble-paired I:C base pair in the unbound state (60). Contrary to this, I-MM bound PU.1 with indistinguishable affinity as MM, suggesting that negligible additional conformational entropy is lost in the DNA upon PU.1 binding.

**Figure 4.**
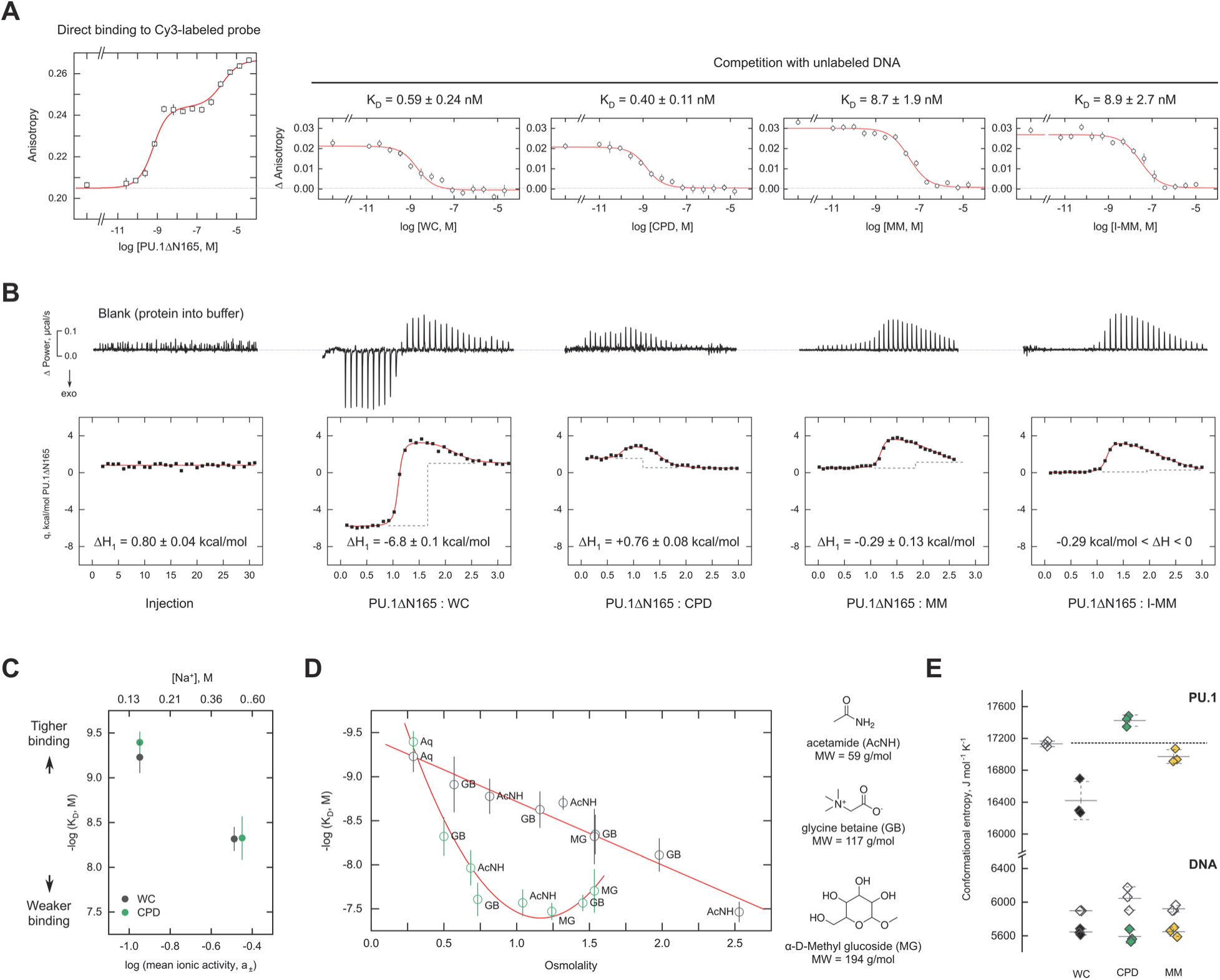
Comparative thermodynamics of ETS interactions with CPD and mismatched DNA. **A**, Equilibrium binding of ΔN165 to the four DNA sequences from the co-crystal structures (WC, CPD, MM, I-MM) as determined by fluorescence polarization. *Cruves* represent fits to the data of an established model in which PU.1 binds sequentially up to a 2:1 complex. Intrinsic affinities to the four sequences were determined by displacement of a PU.1-probe complex at sub-saturating protein concentrations. *Points* represent mean anisotropies ± S.D. of triplicate experiments. **B**, Isothermal titration calorimetry titrations at 25°C. *Curves* represent fits of the same sequential binding model to the thermal data. The change in enthalpy (Δ*H*; long dashes) is computed relative to the apparent heat of diluting ΔN165 into buffer. Bounds for Δ*H* are given in the case of I-MM because the fitted value is smaller than the error in the estimate. **C and D**, Dependence of PU.1 binding to CPD and WC on mean ionic activity (NaCl) or osmolality (non-ionic co-solutes). *Points* are means ± S.D. of triplicate determinations of the equilibrium dissociation constant *K*_D_. The data points at 0.5 M Na^+^ are offset horizontally by 0.5 units to clear their overlap. Osmotic pressure data is labeled by the osmolyte used (shown to the right) at 0.15 M Na^+^. *Aq*, aqueous (no osmolyte added). The spline fit of the CPD data is intended only to guide the eye. **E**, Conformational entropies of free and bound PU.1 and DNA from explicit-solvent MD simulations. Each *point* represents an estimate from the final 5 ns of an independent trajectory.

To directly evaluate the thermodynamic basis of the strong CPD binding by PU.1, we measured the enthalpy change (Δ*H*) of binding by isothermal titration calorimetry [**Figure 4B**]. As detailed in previous analyses (43), the molar heat detected in the initial phase of a protein-into-DNA titration represents the Δ*H* of forming the 1:1 PU.1/DNA complex as captured in the co-crystal structures. In sharp contrast to the **WC** complex, which was markedly exothermic (ΔH ≪ 0) at 25°C, formation of the **CPD** complex was endothermic (Δ*H* > 0), indicating that binding of the CPD target was essentially entropy-driven. Enthalpy contributions to the **MM** and **I-MM** complexes were favorable but quantitatively minor. As published Δ*H* values of native PU.1/DNA complexes track with binding affinity (43), the **CPD** complex represents the first example of high-affinity PU.1 interactions without favorable enthalpic contributions from the noncovalent contacts.

The lack of favorable enthalpic contributions without a corresponding penalty in affinity (free energy change) for CPD binding led us to hypothesize that CPD formation involved significant disposition of solution components for the unbound states. The thermodynamic uptake or release of water, ions, and other low-molecular weight species upon protein-DNA binding is associated with significant change in translational entropy (61). To target these dispositions, we probed the linkage of the equilibrium dissociation constant to the thermodynamic activity of salt and non-ionic osmolytes 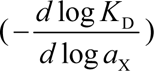. Increased ionic activity ( *a_x_* = *a*_+_ ), as exerted by NaCl, identically decreased binding affinity by ∼10-fold from 0.15 to 0.5 M NaCl for CPD and WC, respectively, indicating no net difference in the disposition of ions from electrostatic contacts between the two complexes [**Figure 4C**].

To target the disposition of non-ionic co-solutes, we measured the impact of glycine betaine, acetamide, and α-methyl glucoside on PU.1/DNA binding. The use of physicochemically dissimilar osmolytes provides a thermodynamic diagnosis of the disposition of co-solutes and water, which is solute-dependent in the former case but colligative in the latter. Osmolyte activity was measured by vapor pressure osmometry and expressed as solution osmolality. In stark contrast with NaCl, non-electrolytes strongly differentiated binding to CPD and WC [**Figure 4D**]. Whereas the binding free energy (which is proportional to − log *K*_D_ ) to the high-affinity WC sequence decreased linearly up to 2.5 osmolal, binding to CPD exhibited a markedly convex dependence. The effects of these non-electrolytes differed from the ionic perturbations at equivalent osmolality (0.97 osmolal for 0.5 M NaCl). Together, these features strongly suggest that the osmolytes are probing general differences in solvent exposure between the two complexes.

The linear and colligative dependence of PU.1 binding to WC sequences on osmolality has been modeled in terms of the change in molecular hydration on binding (44,62–64):

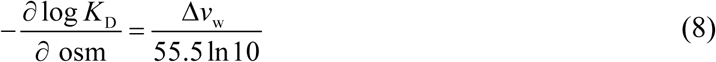

where the coefficient Δ*v*_w_ measures the hydration change ( Δ*v*_w_ > 0 denoting net water uptake) and 55.5 represents the molal volume of pure water. Against this backdrop, the sharp reduction in CPD binding affinity at low osmolality, taken together with the measured positive Δ*H*, is most consistently interpreted as the net release of osmolyte upon complex formation. The convex dependence may be accommodated by a more general formulation of Eq. (8) as the coupled disposition of hydration water and osmolyte (65):

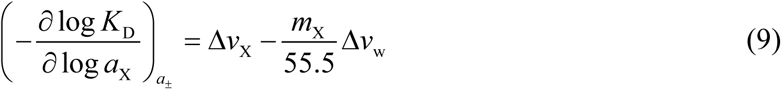

where *m*_X_ is osmolyte molality. Since solute activity coefficient γ_X_ = *a*_X_ / *m*_X_ tracks the osmotic coefficient ϕ_X_ = osm_X_ / *m*_X_ monotonically for the three osmolytes within the molality range of the experiment, the observed convexity in 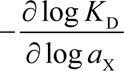 by Eq. (9) indicated a coupled release of osmolyte ( Δ*v*_X_ < 0 ) and water ( Δ*v*_w_ < 0 ).

For a structural interpretation, Parsegian and coworkers have proposed that curvilinear dependence of binding affinity on osmolality represents structural deformation by increasing osmotic pressure (66). As susceptibility to deformation implies a more conformationally dynamic structure, we explore this possibility by molecular dynamics (MD) simulations in explicit solvent. We simulated CPD by restraining the separation distance and/or dihedral of the C5-C6 bonds (see *Materials and Methods*) and estimated absolute conformational entropies from converged MD trajectories of the free and complexed protein and DNA using Schlitteŕs formula (49) [**Figure S3A and 4E**]. Among the DNA duplexes, CPD is associated with the largest loss in absolute entropy upon protein binding, lending credence to this approach. For the protein, CPD-bound PU.1 is uniquely associated with increased conformational entropies relative to the free state. Binding to CPD thus induced excess dynamics in the protein selectively within the complex. In summary, calorimetric characterizations, linkage thermodynamics, and simulations captured a distinct binding mode for CPD that is less rigid, more dynamic, and more deformable than the WC complex within the canonical framework of sequence-specific ETS/DNA structure.

### Dynamic conformational predisposition to CPD formation by the ETS domain

In addition to providing structural insight into binding by ETS proteins to existing CPD, the co-crystal PU.1 complexes also offered a window into how ETS binding predisposes native DNA to CPD formation. In general, duplex dipyrimidine (YY) steps become predisposed to CPD formation as the nucleobases approach the geometry of the DNA photoproduct. More precisely, the probability of reaction (quantum yield) may be modeled in terms of the distance between the C5-C6 bonds and the dihedral angle defined by the two bonds, denoted C5-C6-C6*-C5* (67). ETS proteins such as Ets-1 conformationally align the C5-C6 bonds and shorten their separation, both of which features are associated with enhanced CPD reactivity (21). Here, co-crystal PU.1 complexes also biased the conformation of the corresponding CY base step as in the CPD structure, particularly with respect to bond separation, as did the **MM** and **I-MM** complexes [**Figure 5A**]. To probe the effect of ETS binding on the conformational dynamics at the base step independently of crystal packing, we analyzed equilibrated MD trajectories of the WC sequence, with and without PU.1. The structures in explicit solvent showed that PU.1 binding restricted the distance separation of the C5-C6 bonds into a narrow range near 3.5 Å [**Figure 5B**]. Thus, the dynamic induction of DNA to geometry susceptible to CPD formation also appeared to be a class effect of ETS proteins, in addition to engagement of cognate sites with a formed CPD.

**Figure 5.**
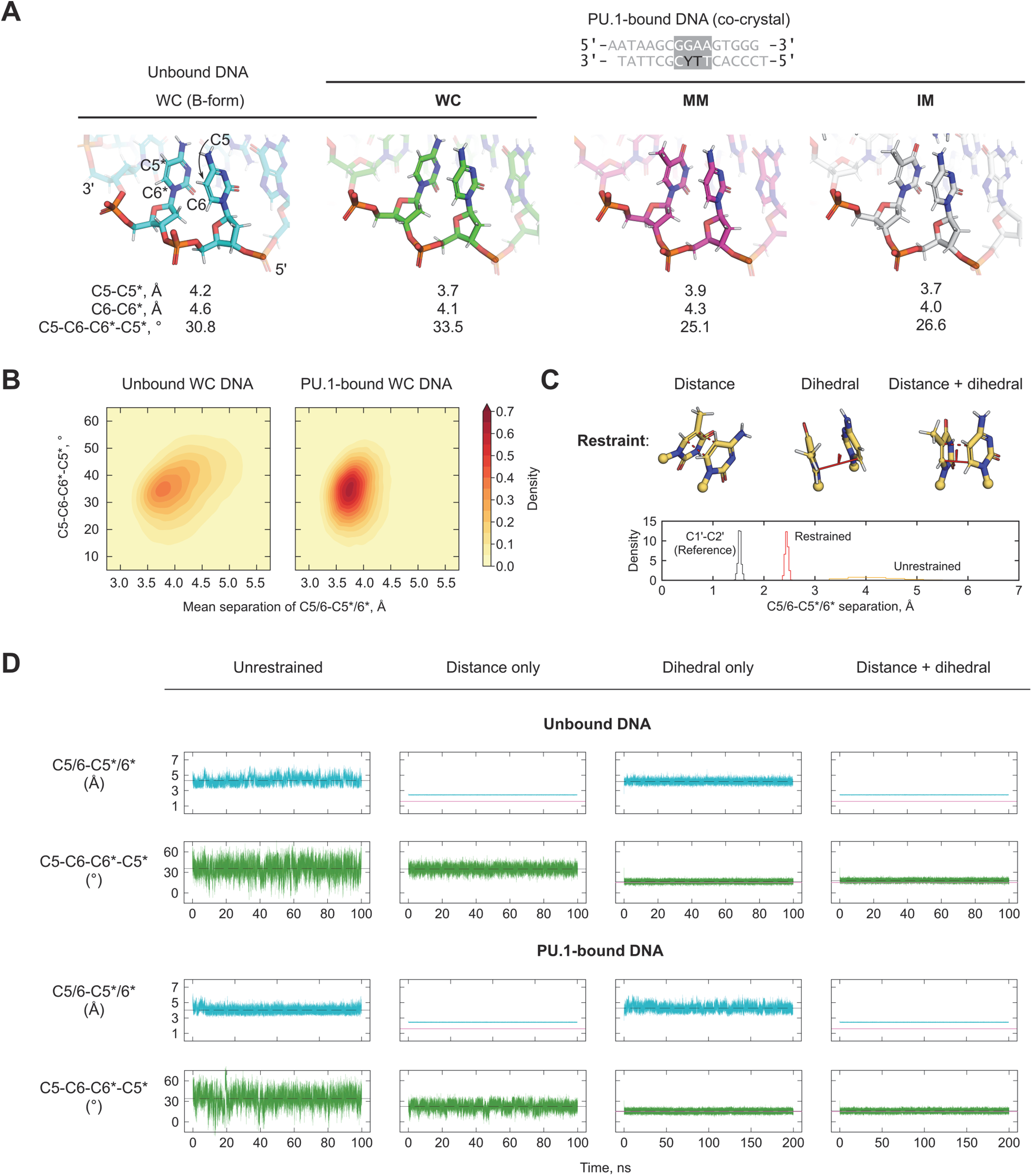
Dissecting the dynamic selection of CPD-reactive conformations at the YY base step by the ETS domain. **A**, Distance and dihedral geometry of the YY step corresponding to CPD among the crystal structures, as defined by the respective C5 and C6 atoms, relative to that for a computed B-form sequence. **B**, Joint disance and dihedral distributions of YY step from explicit-atom MD simulations of the unbound and PU.1-bound WC sequence from the final 100 ns of equilibrated simulations. **C**, Types of restrained MD simulations aimed at dissecting the recognition of CPD-reactive conformation by PU.1. Solid and dashed lines in the histogram refer to the C5-C5* and C6-C6* distances, respectively. The magnitude of the harmonic restrains was calibrated against a “typical” covalent C-C bond as exemplied by C1’-C2’ in deoxyribose. **D**, Distance and dihedral fluctuations of unrestrained and restrained simulations. Dashed and pink lines denote the mean and target value of restrained parameters. Note that distance restraint alone is sufficient for PU.1 to bias dihedral alignment, but not *vice versa*.

While separation of the C5-C6 bonds represented a major conformational perturbation at the CY base step, the quantum yield for CPD also depends strongly on the dihedral alignment of the two bonds (67). Specifically, at separation significantly farther than the C5-C5* and C6-C6* bond lengths of the CPD photoproduct *i.e.,* 1.6 Å for a TpT *cis-syn* dimer (68), the CPD quantum yield decays markedly with the C5-C6-C6*-C5* dihedral angle (67). To reconcile these observations, we hypothesized that CPD-reactive dihedrals are dynamically sampled in a complex conformational landscape. We tested this hypothesis by carrying out restrained MD simulations targeting the separation distance and/or dihedral of the C5-C6 bonds to bias the sampling of conformations [**Figure 5C**]. Harmonic restraints were incorporated with target distance and dihedral as determined quantum mechanically for *cis-syn* TpT CPD (1.6 Å, 15°) (68) and force constants were tuned to reproduce bond-distance dynamics matching typical covalent C-C bonds in the structure (chosen as C1’-C2’ of deoxyribose) [**Figure S3B**]. Under these specifications, the time-averaged restrained separation and dihedral between the C5-C6 bonds were 2.5 Å and 16°, compared with 4.3 Å and 36° in unrestrained controls.

The distance and dihedral dynamics in equilibrated structures of the WC sequence were compared alone or in complex with the ETS domain of PU.1 [**Figure 5D**]. For the unbound DNA, imposition of either distance and/or dihedral restraints reduced the fluctuations of the free parameters but did not significantly alter its mean magnitude. In contrast, PU.1-binding significantly decreased C5-C6-C6*-C5* to 23° (a reduction of ∼40%) when *only* distance restraints were applied. This effect was not reciprocal, however, as dihedral restraints alone showed little effect on the C5-C6 distance. The simulations thus suggested that predisposition by ETS binding is conformationally funneled through the capture of dihedral-aligned states by shortening C5-C6 separation.

## DISCUSSION

DNA binding sites of ETS family TFs are among the most highly mutated genomic regions in skin cancers, such as melanoma (10,13). Previous studies have suggested that recurrent mutations at these sites are the result not only of ETS binding stimulating UV-induced CPD formation, but also because ETS binding inhibits subsequent repair of these CPD hotspots (14,21,22,27). However, the molecular mechanism by which ETS family members persistently bind damaged sites containing CPDs and restrict their repair was previously unclear. Here, we show that paralogs from each of the three different ETS classes (i.e., ETS1, ELF1, and PU.1) bind with high affinity to cognate DNA sites containing a central TpT CPD, often with higher affinity than undamaged sites. Furthermore, we show that DNA binding by each of the ETS paralogs can inhibit repair of the CPD lesion by restricting its access to a model repair enzyme. We solved the structures of the ETS family member PU.1 bound to CPD-containing and mismatched DNA sites, and characterized the thermodynamic binding profile by which PU.1 binds a damaged DNA site with high affinity. These findings provide a molecular rationale for how ETS family members accommodate a CPD lesion in their binding site to prevent access by repair enzymes, and thereby promote mutation hotspots in melanoma and other skin cancers.

Environmental UV radiation penetrates through the epidermis to the dermal layer of the skin to an extent dependent on wavelength and skin-related factors (69). These layers contain cells, such as keratinocytes and fibroblasts that are responsible for the structural components of skin, as well as melanocytes and immune cells (*e.g.*, Langerhans cells) with specialized functions in the viable epidermis. Members from Class I to III of the ETS superfamily are broadly expressed in these tissues (30), either as part of normal skin function and turnover, or in specific pathophysiology (31). Using the *DPH3* promoter as a typical model of UV-damaged ETS target, the observations that ETS factors from all three classes can bind CPD-containing sites comparably as well as native cognate DNA sites, and that ETS binding competes with DNA repair enzymes, suggest CPD lesions are subject to repair inhibition by the ETS family as a whole. As the tested ETS relatives are highly representative of their respective classes in DNA selectivity on the one hand (51), but also derive from highly diverse domain context in the full-length protein (29), the data supports the proposal of high-affinity CPD engagement as a novel class property of DNA-binding ETS domain.

Although the Class III member PU.1 binds the native DPH promoter less tightly than ETS1 and ELF1, as expected from motif preference, the presence of a centered CPD nevertheless stimulated binding (Figure 1C) and cleavage inhibition (Figure 2C) on par with the other paralogs. While the co-crystallographic evidence showed that the gross architecture and base-specific DNA contacts within the consensus remain similar to the WC complex (Figure 3B and C), the thermodynamics (Figure 4) unveiled a wholly different binding mode for CPD. Previously, entropy-only driven binding of ETS factors was only observed for low-affinity to suboptimal WC-matched binding sites (43,62). Experimental agreement of the linkage thermodynamics (63) with osmotic stress (70) has led to a model in which PU.1 binding specificity is contingent on the extent of uptake of hydration water (44,64). Here, CPD binding by PU.1 is also entropy-driven only but proceeds with no penalty in free energy (affinity). Accordingly, the linkage thermodynamics with osmolytes revealed a distinct profile consistent with a less rigid (more deformable) structure. The differential structural perturbations between the WC and CPD complexes involve in part disordered structures which are not resolved crystallographically and attributed by all-atom simulations to the CPD-bound protein in terms of increased conformational entropy.

A high-throughput binding study has previously shown that the presence of UV damage alters the sequence selectivity of ETS1 and ELK1 (20). This study found that for UV-irradiated DNA, ETS1 and ELK1 favor binding to 5’-ATCC-3’ sequences instead of sites that contain a 5’-TTCC-3’ consensus. Our binding data provide a molecular rationale for this change in sequence specificity, as a site-specific CPD in the variable TpT dinucleotide in the ETS binding consensus (i.e., **CPD_-2_**, 5’-TT<u>CC</u>-3’) reduces the binding affinity by each of the ETS paralogs. In contrast, 5’-ATCC-3’ sites are likely favored following UV irradiation because they are unable to form this low affinity lesion, but can form the high affinity CPD at the TpC dipyrimidine (i.e., **CPD_-1_**). ETS1 and ELK1 are both Class I factors. The stimulated binding of PU.1 and ELF1 by a consensus-centered **CPD_-1_**, but decreased affinity for **CPD_-2_**, suggests that the same alteration of specificity may occur with other classes of ETS factors. Structurally, the centered CPD (i.e., **CPD_-1_**) is a non-contacted moiety in the consensus, which indicates that its effects on binding involves a change in indirect readout. The helical contortion (Figure 3G) that ultimately presents in a formed CPD is not accessible to the double helix prior to photodamage. As our MD simulations show that shortening of the base step, not dihedral alignment, represents the conformational predicate to CPD formation by UV radiation, a reasonable suggestion is that other base-adjacent photoproducts at the same position, such as the pyrimidine-pyrimidone (6–4) photoproduct, may also exhibit enhanced affinity to ETS factors as well.

## CONCLUSION

In conclusion, these findings provide a structural and thermodynamic explanation for how ETS family proteins can accommodate a helix-distorting DNA lesion in their binding sites. Our findings that ETS TFs bind with high affinity (in some cases even higher than an undamaged site) to a deaminated CPD in the center of the ETS binding consensus can explain cellular results indicating that deaminated CPDs accumulate at this relative location in ETS binding sites and why repair of these CPDs is inhibited in cells. Importantly, this position also precisely coincides with the relative location of mutation hotspots in melanoma (9,21–27), many of which may play an important role in melanomagenesis. Furthermore, our discovery that ETS binding affinity significantly varies with the location of the CPD lesion in the binding motif can explain a prior study indicating that UV damage alters ETS sequence specificity (20). The stimulated recruitment of PU.1 to suitably positioned CPDs at ETS motifs that are not preferred targets in their undamaged state further suggests that the pool of ETS-based inhibitors of DNA repair may exceed the complement of ETS factors that bind the undamaged site. As many other TFs show altered sequence-specificity in UV-damaged DNA (20), we hypothesize that a mechanism similar to that described here may regulate the binding of other TFs to damaged DNA sites, including many TFs (*e.g.,* c-MYC, EGR1, and p53) that play critical roles in carcinogenesis.

## ACKNOWLEDGEMENT

We thank the beamline staff of the National Synchrotron Light Source II (NSLS-II) at Brookhaven National Laboratory (Upton, NY) for their support during the X-ray data collections. We acknowledge Dr. Amanda V. Albrecht for technical assistance. This investigation was supported by NIH grants HL155178 and TR004842 (to G.M.K.P.) and ES028698, ES032814, ES035139, and ES035888 (to J.J.W.) and NSF grant MCB 2028902 (to G.M.K.P and M.W.G).

## DATA AVAILABILITY STATEMENT

Coordinates and validation reports for the **CPD**, **MM**, and **I-MM** structures have been deposited with the RCSB PDB under accession codes 9YNZ, 9OA4, and 9OB0. Gel images and all other relevant data are available from the corresponding authors upon reasonable request.

## AUTHOR CONTRIBUTIONS

Conceptualization, S.S., J.J.W., G.M.K.P.; Methodology, S.S., J.R.T., M.W.G., G.M.K.P.; Investigation, S.S., J.R.T., M.W.G., M.F.L., S.P.A., M.E.E., P.J.H., G.M.K.P.; Formal data analysis, S.S., J.R.T., M.W.G., M.F.L., G.M.K.P.; Writing and editing, S.S., J.R.T., M.E.E., M.W.G., P.J.H. J.J.W., G.M.K.P.; Data visualization, S.S., J.J.W., G.M.K.P.; Supervision, J.J.W., G.M.K.P; Funding acquisition, M.W.G., J.J.W., G.M.K.P.

