## Supplemental Table S1 and Figures S1 to S3 for "Molecular basis of UV lesion binding and repair inhibition by ETS-family transcription factors"


January 13, 2026


[1] School of Molecular Biosciences, Washington State University, Pullman WA 99164

[2] Department of Chemistry, Georgia State University, Atlanta, GA 30303

[3] Department of Biology, Georgia State University, Atlanta, GA 30303

[4] Department of Chemistry, University of Idaho, Moscow, ID 83844

[#] Present address: Department of Molecular Biosciences, University of South Florida, Tampa, FL 33620

[##] Present address: U.S. Army Medical Research Institute of Chemical Defense, Aberdeen Proving Ground, MD 21010

[###] Present address: Department of Pharmacology, University of Michigan Medical School, Ann Arbor, MI 48109

[*] To whom correspondence should be addressed at: Biotechnology Life Sciences 241, Pullman, WA 99164-7520, U.S.A. (J.J.W.) or P.O. Box 3965, Atlanta, GA 30302-3965, U.S.A. (G.M.K.P.)..

**SUPPLEMENTAL METHODS**

*Synthesis of CPD-containing oligonucleotides*. CPD-modified oligonucleotides were synthesized via machine-assisted solid-phase DNA synthesis (0.2 µmol scale, 500 Å LCAA-CPG solid support). Standard protocols were used for incorporation of conventional nucleobase-protected DNA monomers (DMT-on). The *cis-syn* thymine dimer phosphoramidite (Glen Research) was incorporated via hand-coupling (anhydrous acetonitrile, 4,5-dicyanoimidazole, 15 min coupling time) in low light. Following the incorporation of the final monomer, the methyl phosphate protecting groups of the dimer were removed by treating the oligo-bound solid support with thiophenol/triethylamine/THF (1:2:2 v/v/v) for 45 min at room temperature in low light. The solid support was subsequently washed with THF (1×), methanol (5×), and acetonitrile (3×), and dried with argon flow for 5 min. The CPD oligos were treated with 32% aqueous ammonia (55°C, 17 h) to remove nucleobase protecting groups and cleave the oligos from the solid support. The ammonia solution was evaporated off using a centrifugal vacuum concentrator and the crude CPD oligos were resuspended in 500 µL HPLC water. Detritylation and purification of CPD oligos was performed at room temperature using TOP-DNA 150 mg oligonucleotide cartridges (Agilent) following the manufacturer's recommendation. Thus, TOP-DNA cartridges were secured in a vacuum manifold, conditioned with 0.5 mL HPLC-grade MeCN (flow rate of ~2 drops/s), and equilibrated with 1 mL of 2 M aqueous triethyl ammonium acetate (TEAA). An aqueous NaCl solution (100 mg/mL, 500 µL) was added to the resuspended CPD oligos and the resulting mixture loaded on the cartridge. The cartridge was rinsed with an additional 2 mL of the NaCl solution to elute impurities. The CPD oligos were then detritylated (2 mL, 5% aq. $CF_3COOH$), rinsed with 2 mL of HPLC-grade water, and eluted from the cartridge (0.5 mL, 50:50 MeCN:H2O, v/v). The

CPD oligos were dried using a centrifugal vacuum concentrator and reconstituted in HPLC-grade water. The purity and identity of the synthesized CPD oligos was established using analytical HPLC (XTerra MS C18 column: 0.05 M TEAA and MeCN gradient) and LC-ESI-MS analysis (Waters Acquity C18 column; TEAA and MeCN gradient) recorded on a quadrupole time-of-flight (Q-TOF) mass spectrometer. The raw signals were deconvoluted using the Max Ent software provided with the spectrometer to obtain molecular ion peaks.

*NMR characterization of CPD-containing and mismatch duplexes*. $^{1}$H NMR spectra were collected on a Bruker Avance III HD 600 MHz NMR equipped with a 5 mm TXI probe. DNA duplex samples were prepared in 10 mM $K_2HPO_4$/$KH_2PO_4$, 50 mM KCl, 1 mM EDTA, and 10% $D_2O$. Samples were adjusted to pH 6.5. 1D $^{1}$H imino spectra were acquired using 1-1 jump-return solvent suppression at 293 K. 1-1 NOESY spectra were collected (2048 × 800) at 293 K using 150 ms mixing time. Chemical shifts assignments were initially made using a fully WC-paired construct from the NOESY spectrum and 1D spectra acquired at 278 to 323 K. These assignments were then used to assign the CPD and MM duplexes by analogy.

*Enzymatic validation of CPD-containing oligonucleotide*. To confirm the presence of a CPD in the single-stranded oligonucleotide substrate used for X-ray crystallography studies, a hybridization reaction was performed with the 16-nucleotide (nt) damage-containing substrate ("CPD oligo", 5′-TCCCACTT^TCGCTTAT-3′), a 9-nt 3′ 6-FAM labeled oligonucleotide (5′-CAGATCAAC-3′-6-FAM), and a 25-nt oligonucleotide complementary to the other fragments ("Bottom oligo", 5′-GTTGATCTGATAAGCGAAAGTGGGA-3′). Approximately equimolar concentrations of the CPD-containing oligonucleotide and the bottom oligo were used to assemble the hybridization reaction; however, half of this concentration was used for the 6-FAM oligo in order to minimize the amount of signal on the gel derived from unligated 6-FAM-containing oligonucleotide. The reactions were assembled in 1× T4 ligase buffer (New England Biolab) and run in a thermocycler that performed an initial incubation at 95ºC for 3-5 minutes, followed by decrease in temperature at a rate of 1ºC per minute for a total of 70 cycles.

Following the hybridization reaction, T4 DNA ligase (NEB) was added and the reaction was incubated overnight at 16°C. The following day, the reaction extracted in 25:24:1 phenol:chloroform:isoamyl alcohol (Fisher) and ethanol-precipitated with added glycogen for several hours at -20°C prior to centrifugation. Pellets were resuspended in water, half of which was saved as a control, and the other half was digested with T4 PDG (NEB) in the manufacturer's supplied buffer (1×) at 37°C for one hour. Prior to electrophoresis, samples were equilibrated in 0.15 N NaOH and heated at 95ºC to promote cleavage of the DNA at abasic sites rendered from cleavage of the CPD damage. Finally, 2.5 volumes of formamide (Millipore Sigma) were added to the samples, which were heated again at 95ºC to promote denaturation of secondary DNA structure.

Electrophoresis of the digested sample (along with controls including the 6-FAM oligo alone, the hybridized but unligated reaction, and the ligation reaction prior to T4 PDG digest) was performed in preheated denaturing (6.1 M urea) 12% polyacrylamide gels in 1× TBE buffer. Gels were imaged on a GE Typhoon using the setting for FAM-labeled substrates and images were visualized with ImageQuant TL.

As an alternate method of substrate confirmation, the CPD-containing oligo was hybridized to the complementary 'Bottom oligo' in water or 0.1× TE pH 7.5 in a thermocycler as described above, and then digested directly with T4 PDG in the supplied buffer for 2 h at 37°C. A control reaction was also performed using an oligo identical in sequence to the CPD-containing oligo but lacking the CPD lesion (*i.e.,* “No CPD oligo”, 5′-TCCCACTTTCGCTTAT-3′). After digestion, samples were heated in denaturing conditions similar to the method described above, but with slightly shorter incubation times (~9 min). Samples were electrophoresed as described above. Following electrophoresis, the gel was stained in 1x SYBR Gold (Invitrogen), imaged on a GE Typhoon, and images were visualized with ImageQuant TL.

**Table S1. Refinement statistics of co-crystallographic models**

| **Protein** | hPU.1(165-270) | hPU.1(165-270) | hPU.1(165-270) |
|---|---|---|---|
| **DNA** | CPD | MM | I-MM |
| **PDB ID** | **9YNZ** | **9OA4** | **9OB0** |
| Wavelength | 0.9201 | 0.9201 | 0.9201 |
| Resolution range | 23.85 - 2.05 (2.123 - 2.05) | 24.19 - 1.42 (1.471 - 1.42) | 24.17 - 1.79 (1.854 - 1.79) |
| Space group | P 1 21 1 | P 1 21 1 | P 1 21 1 |
| Unit cell | 42.966 58.879 45.925 90 117.593 90 | 43.134 60.695 44.802 0 116.593 90 | 42.996 60.588 44.909 0 116.789 90 |
| Total reflections | 47521 (4874) | 166740 (16873) | 75493 (7631) |
| Unique reflections | 12776 (1256) | 38841 (3882) | 19169 (1873) |
| Multiplicity | 3.7 (3.9) | 4.3 (4.3) | 3.9 (4.1) |
| Completeness (%) | 99.02 (99.52) | 99.26 (99.67) | 98.19 (97.24) |
| Mean I/sigma(I) | 7.43 (1.92) | 16.66 (2.89) | 10.80 (2.41) |
| Wilson B-factor | 35.53 | 20.01 | 24.85 |
| R-merge | 0.105 (0.4288) | 0.04254 (0.428) | 0.07295 (0.5132) |
| R-meas | 0.122 (0.4957) | 0.04924 (0.4867) | 0.08427 (0.5901) |
| R-pim | 0.06139 (0.2459) | 0.02424 (0.2282) | 0.04171 (0.2885) |
| CC1/2 | 0.993 (0.915) | 0.997 (0.926) | 0.998 (0.857) |
| CC* | 0.998 (0.978) | 0.999 (0.981) | 0.999 (0.961) |
| Reflections used in refinement | 12776 (1253) | 38841 (3874) | 19169 (1870) |
| Reflections used for R-free | 1318 (132) | 1985 (201) | 1952 (194) |
| R-work | 0.2063 (0.2776) | 0.1592 (0.2168) | 0.1657 (0.2391) |
| R-free | 0.2440 (0.3439) | 0.1946 (0.2516) | 0.2106 (0.2890) |
| CC(work) | 0.961 (0.895) | 0.938 (0.953) | 0.971 (0.913) |
| CC(free) | 0.934 (0.724) | 0.936 (0.929) | 0.969 (0.865) |
| Number of non-hydrogen atoms | 1445 | 1639 | 1587 |
| macromolecules | 1334 | 1400 | 1378 |
| ligands | 64 | 0 | 31 |
| solvent | 71 | 239 | 188 |
| Protein residues | 91 | 91 | 91 |
| RMS(bonds) | 0.013 | 0.008 | 0.012 |
| RMS(angles) | 1.49 | 1.04 | 1.44 |
| Ramachandran favored (%) | 98.88 | 97.75 | 98.88 |
| Ramachandran allowed (%) | 1.12 | 2.25 | 1.12 |
| Ramachandran outliers (%) | 0 | 0 | 0 |
| Rotamer outliers (%) | 5.19 | 0 | 0 |
| Clashscore | 4.04 | 1.57 | 1.58 |
| Average B-factor | 45.82 | 29.39 | 28.51 |
| macromolecules | 41.21 | 26.62 | 26.12 |
| ligands | 48.05 | - | 32.98 |
| solvent | 44.44 | 39.54 | 37.38 |
| Beamline | 17-ID-1 AMX | 17-ID-1 AMX | 17-ID-1 AMX |
| Detector | Dectris Eiger 9M | Dectris Eiger 9M | Dectris Eiger 9M |
| Oscillation Angle | 0.1° | 0.1° | 0.1° |
| Frames Collected | 2000 | 2200 | 2000 |

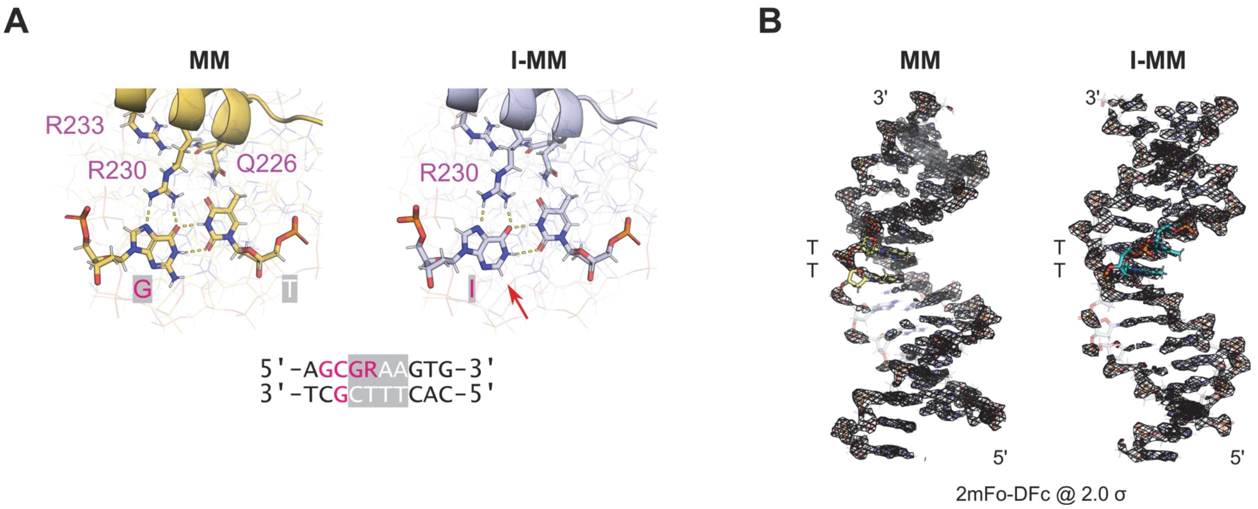


**Figure S1. Co-crystallographic complex of the ETS domain of PU.1 with inosine-substituted DNA mismatch (I-MM)**. **A**, Protein/DNA interface at the major groove consensus of the refined structures. Arrow points to the 2-deaminated position, which is non-contacted in **I-MM**, at the wobble base pair. **B**, Electron density (2mFo-DFc) maps of the PU.1-bound DNA. Note the similar spatial distributions along the DNA sequence.

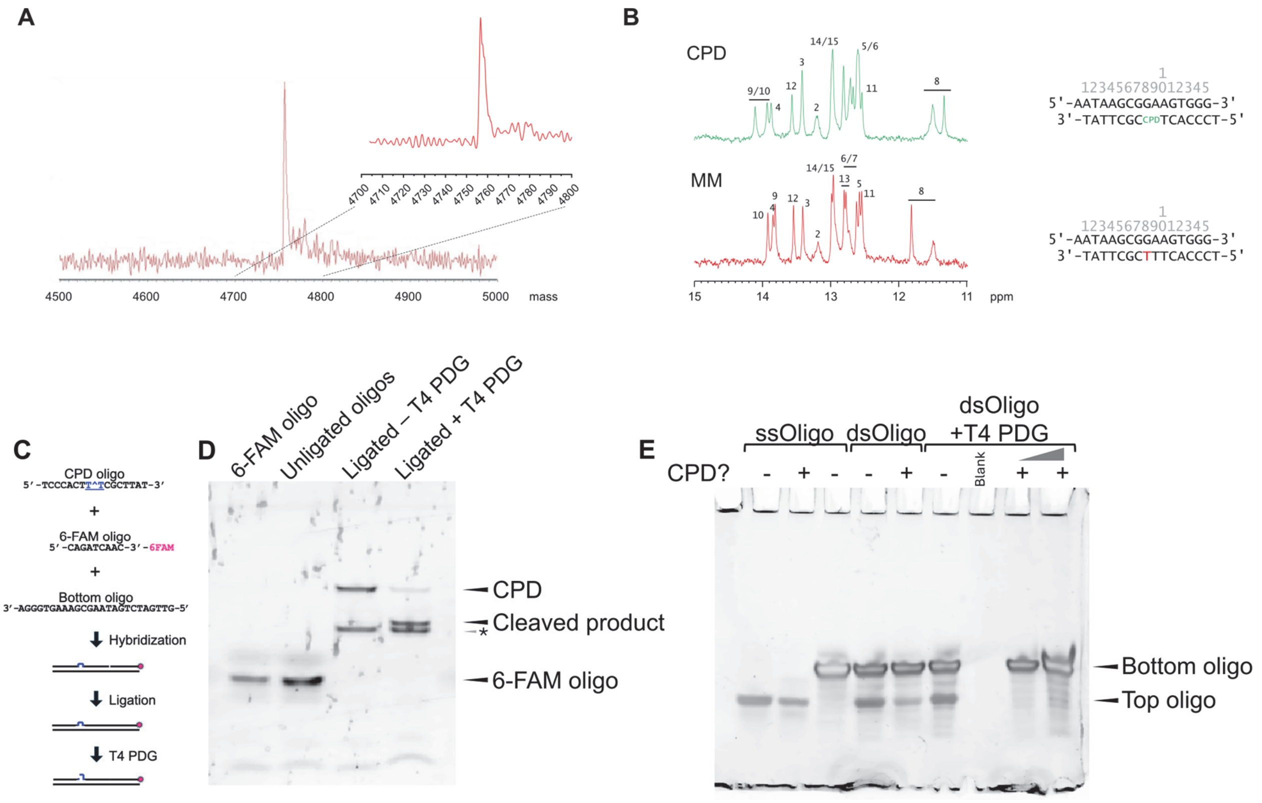


**Figure S2. Confirmation of the CPD substrate used in the co-crystal structure with PU.1**. **A**, Deconvoluted ESI-MS spectrum of the CPD-bearing strand following HPLC purification. The ESI solvent was 50 mM tetraammonium acetate in acetonitrile. **B**, Imino $^1$H NMR of the CPD and MM sequence in duplex with the same complementary oligo as WC in 50 mM $K_2HPO_4$/$KH_2PO_4$ (pH 6.5) containing 1 mM EDTA at 293 K. **C**, Experimental design for ligating CPD-containing oligonucleotide (CPD oligo) used for X-ray crystallography experiments to a small oligonucleotide with a 3′ 6-FAM fluorescent moiety (6-FAM oligo), so that CPD cleavage by T4 PDG could be monitored by fluorescence imaging. **D**, Denaturing PAGE analysis of CPD-containing oligonucleotide used in the co-crystal structure in the presence or absence of T4 PDG. A 9 nt 6-FAM labeled oligonucleotide (run alone, first lane) was hybridized to a second 25 nt oligo ('Bottom oligo') that contained regions complementary to it as well as the 16 nt CPD-containing oligo (immediately upstream, unligated hybridization reaction run in second lane). Once hybridized, the DNA was treated with T4 ligase to ligate the 6-FAM oligo with the CPD oligo (third lane, top band). The hybridized, ligated dsDNA was then digested with T4 PDG, which cleaves CPDs (fourth lane). A recurrent band of undetermined origin (labeled with *) consistently appeared in the ligation and T4 PDG-treated reactions, which is likely due to degradation or breakage at the site of the CPD, potentially due to excessive handling. **E**, SYBR gold staining of denaturing urea PAGE analysis of T4 PDG digested CPD-containing oligonucleotide used in the co-crystal structure. The single stranded CPD-containing oligonucleotide (16 nt, lane 2) was run alongside an undamaged oligonucleotide of the same sequence (16 nt, lane 1) and a complementary oligonucleotide that contained 9 additional nucleotides of non-complementary sequence ('Bottom oligo', 25 nt total, lane 3). Sequences of CPD and bottom oligos are shown in *Panel C*. A longer complementary oligo was used so it could be distinguished on a SYBR-stained denaturing gel from the CPD-containing oligo, due to the difference in sizes. The undamaged (*i.e.,* No CPD) control and CPD-containing top oligos were independently hybridized to the complementary Bottom oligo (lanes 4 and 5, respectively) and then digested with T4 PDG, which cleaves CPD bases. No cleavage was observed in the hybridized undamaged (no CPD) DNA (lane 6). However, T4 PDG cleavage was evident in the CPD-containing oligo (lanes 8 and 9, 2× volume is loaded in the latter), as the band for the top oligo disappears. No cleavage products were detected, likely because they are too small to be stained effectively by SYBR gold.

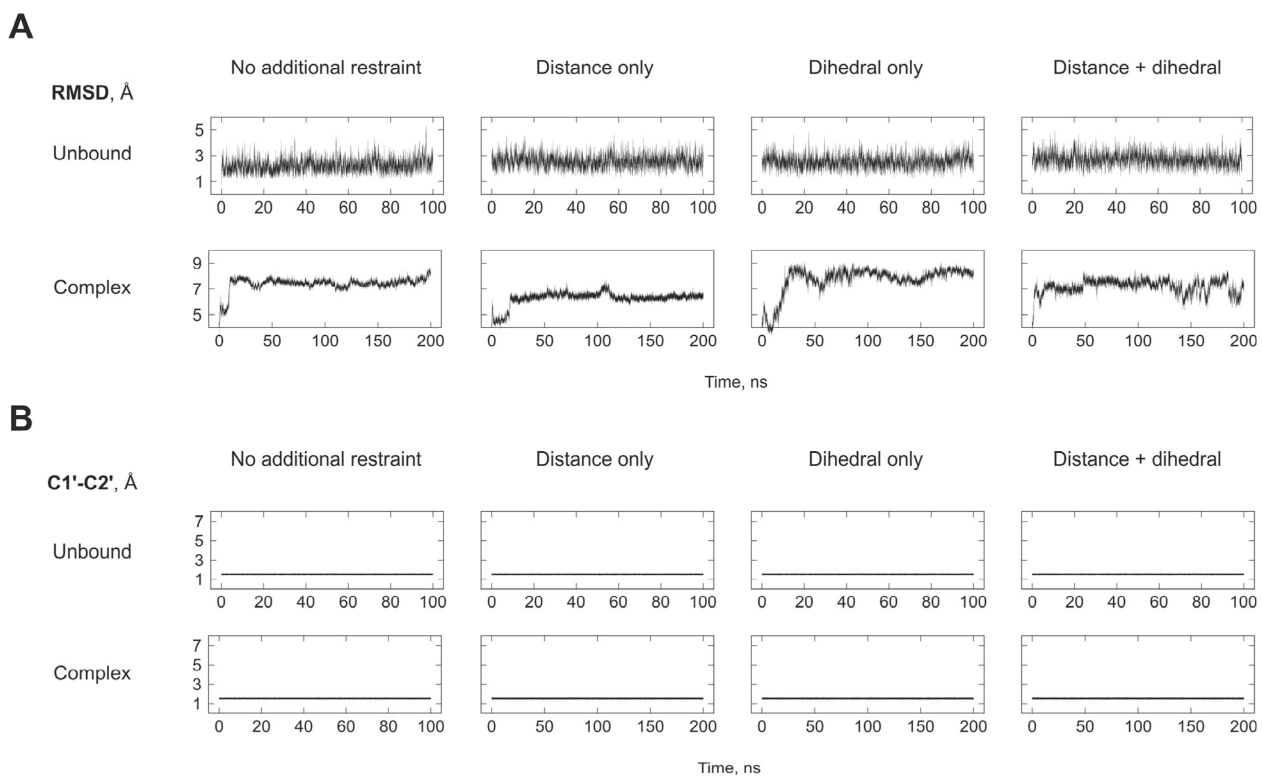


**Figure S3. Equilbration and calibration of restrained molecular dynamics simulations**. **A**, Representative self-versus-self RMS deviation of unbound and PU.1-bound DNA sequences. For complexes, the final 100-ns interval is taken for analysis. **B**, Representative distance fluctuations of the C1´-C2´ bond in deoxyribose, used to calibrate force constants used in the restrained simulations.
